# Scream’s roughness confers a privileged access to the brain during sleep

**DOI:** 10.1101/2022.09.05.506631

**Authors:** G Legendre, M Moyne, J Dominguez-Borras, S Kumar, V Sterpenich, S Schwartz, LH Arnal

## Abstract

During sleep, recognizing threatening signals is crucial to know when to wake up and when to continue vital sleep functions. Screaming is perhaps the most salient and efficient signal for communicating danger at a distance or in conditions of limited visibility. Beyond the intensity or the pitch of the sound, rapid modulations of sound pressure in the so-called roughness range (i.e. 30-150 Hz) are particularly powerful in capturing attention and accelerating reactions. Roughness is an acoustic feature that characterizes alarm signals such as screams. However, whether rough sounds are also processed in a privileged manner during sleep is unknown.

We tested this hypothesis by stimulating sleeping human participants with low-intensity screams and neutral calls. We found that screams trigger more reliable and better time-locked responses in wakefulness and NREM sleep. In addition, screams boosted sleep spindles, suggesting elevated stimulus salience. The increase in sleep spindle power was linearly proportional to the roughness of vocalizations, but not to their pitch.

These findings demonstrate that, even at low sound intensity, scream’s roughness conveys stimulus relevance and enhances processing in both the waking and sleeping states. Preserved differential neural responses based on stimulus salience may ensure adaptive reactions –and ultimately survival– in a state where the brain is mostly disconnected from external inputs.

## INTRODUCTION

The ability to detect threat cues regardless of current attentional or vigilance state is essential to react to danger and to ensure survival. During sleep, it is crucial to detect threatening signals and return to wakefulness to respond to imminent danger. Some sounds have the capacity to disrupt sleep continuity more than others, especially if they are highly relevant for the sleeper. For instance, parents wake up more often upon hearing the cry of their own baby as compared to that of others (Formby, 1967). Similarly, hearing one’s own name (as compared to someone else’s name) induces a distinctive EEG response during sleep (Beh & Barratt, 1965; Blume et al., 2017; Perrin et al., 1999). Other auditory properties carrying meaningful information for the sleeper such as emotional prosody and familiarity of voices, can be discriminated by the sleeping brain (Blume et al., 2017, 2018; Chen et al., 2016; Moyne et al., 2022). This ability is particularly relevant to detect potential threats during sleep. Auditory emotional salience processing during sleep has been mostly demonstrated by measurements of number of awakenings (Formby, 1967) and event-related potentials (ERPs; Chen et al., 2016; Perrin et al., 1999) in the past. However, changes in EEG power (Blume et al., 2017, 2018), and sleep oscillations (Beh & Barratt, 1965) are also evoked after the detection of salient stimuli. Noteworthy, auditory stimuli played below the awakening threshold can trigger slow-waves (i.e. large deflections of the EEG signal between 1 and 4 Hz) and sleep spindles (i.e. waxing and waning oscillation with a peak frequency between 12.5 and 14 Hz). Evoked slow-waves are traditionally considered markers of unexpectedness (see Colrain, 2005) but as spindles are often following evoked or spontaneous slow-waves, their link with unexpected events is less clear (Fernandez & Lüthi, 2019). Thus, the detection and salience of a stimulus by sleeping participants can be probed through ERPs and the power of meaningful sleep oscillations such as delta and sigma frequency range (i.e. 1-4Hz and 13-16Hz).

Most of the aforementioned studies on auditory emotional processing during sleep were conducted using complex, natural vocalizations as stimuli. However, several acoustic parameters may contribute to the emotionality of vocalization. In particular, recent works focusing on the acoustics of human alarm calls have shown that screams, unlike emotionally neutral vocalizations, exhibit fast amplitude modulations in the so-called roughness range (i.e. 30-150 Hz; Arnal et al., 2015; see Fig. 1B and 1C). Roughness, to a higher extent than pitch or slower speech features (e.g. syllables), is perceived as aversive and is mostly exploited for alarm signaling (whether natural or artificial), thereby ensuring the selective and efficient processing of warning sounds (Arnal et al., 2015). Importantly, roughness in screams induces more accurate and faster reactions than neutral sounds, even at low signal to noise ratio (Arnal et al., 2015). This ensures that alarm vocalizations can be processed efficiently by the brain, even from a distance. Given that scream’s roughness triggers privileged brain responses and reactions at wake, we hypothesized that this feature might induce arousing responses such as slow-waves and sleep spindles in the sleeping brain.

**Figure 1.**
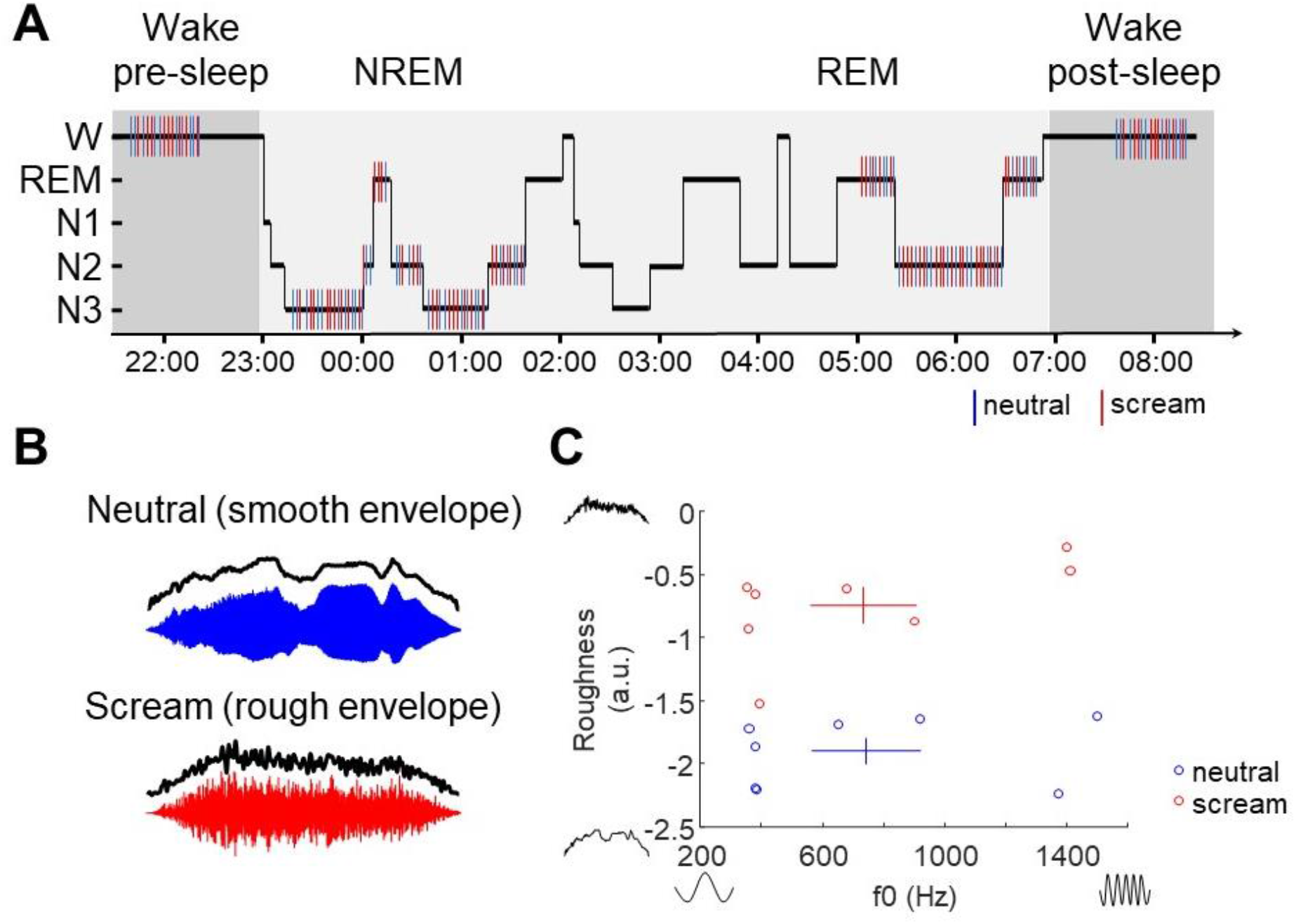
Stimulation protocol and acoustic stimuli characterization. **A**. Example hypnogram of one participant depicting the presentation of stimuli as a function of sleep stages over one night. Stimuli were first played in a brief session of wakefulness before sleep, during the three first hours of sleep, during the two last hours of sleep and after sleep, during another session of wakefulness. **B**. Examples of two auditory stimuli waveforms in each condition: neutral (blue) and screamed (red) vocalizations. Overlaid black line visually depicts that neutral voices are smoother than rough screams. **C**. Eight screams and eight neutral vocalizations (from Arnal et al. 2015) were presented to participants in a pseudo-randomized fashion. This subset of vocalizations was chosen to cover a wide range of fundamental frequencies (f0, paired across neutral and screamed vocalizations) and roughness levels. As quantified in Arnal et al. 2015, the scatter plot reveals that screams and neutral vocalization mainly differ in their roughness levels. Crosses represent the mean (center) and standard error of the mean (tails) of pitch (x-axis) and roughness (y-axis) for screams (red) and neutral vocalizations (blue).

To test this hypothesis, we stimulated participants with screams and neutral vocalizations at a very low sound intensity during wakefulness and sleep. We carefully controlled stimuli so that neutral vocalizations and screams span two main dimensions: roughness and pitch, all other parameters (intensity, attacks) being equal. Recent studies probed the detection during wakefulness and sleep of a hard-to-detect noise using intertrial phase coherence (ITPC) of brain responses in the theta frequency range (Andrillon et al., 2015, 2017) which reflect the degree of similarity of brain response across trials. In addition to traditional ERPs analyses and power analyses of typical frequency ranges such as delta and sigma, we evaluated the ability of the brain to discriminate between neutral vocalizations and screams with ITPC. During wakefulness, we observed more reliable evoked potentials to screams compared to neutral vocalizations. During NREM sleep, we found that the initial auditory response to screams was also more reliable, and that screams induced more spindles than neutral vocalizations. Critically, the increase in spindle activity was linearly related to the roughness of vocalizations, but not to their pitch, indicating a specificity of roughness to enhance the salience of vocalizations.

## MATERIAL AND METHODS

### Participants

Twenty-three healthy participants were recruited via posters displayed at the University of Geneva. Participants reported normal audition, with no known neurological or psychiatric disorder, and no reported phobias or drug consumption. All participants signed an informed consent validated by the local ethical committee for research (Commission cantonale d’éthique de la recherche de Genève). All participants were blind to the experimental conditions but were informed that sensory stimulations would be played during their sleep. The data from six participants were excluded from the analyses, due to poor EEG signal (N=3) or to bad sleep quality (N=3). The data from four other participants were rejected from the analysis due to insufficient number of unartefacted trials (<100 trials) in wakefulness or NREM sleep. Due to the limited number of trials in REM sleep, we did not investigate this sleep stage. The final sample includes 13 participants (8 females; age range = 21.69±1.60 - mean±standard deviation). For more information about participant demographics, see Moyne et al. (2022).

### Stimuli

Two distinct sets of vocalizations were played to the participants during wakefulness and during one night of sleep. The first set consisted of pseudowords and the ensuing results have already been reported elsewhere (Moyne et al., 2022). The second set of stimuli was used in the current experiment and consisted of neutral and screamed meaningless [‘Aaaah’] vocalizations detailed in Arnal et al. (2015). In the Scream condition, 8 meaningless screamed vocalizations were uttered by 8 different actors (4 males and 4 females). In the Neutral condition, 8 loud (but not screamed) vocalizations were produced by the same actors. Neutral vocalizations were pitch-shifted to match the fundamental frequency (F0) of the scream of the same actor. All vocalizations were acquired at 44.1 kHz and subsequently resampled at 16 kHz. All vocalizations were edited to last 750 ms and normalized for sound attacks using onset and offset sine ramping of 100 ms and for intensity (using RMS normalization). The roughness values of each vocalization (Fig. 1B and 1C) was estimated using the Modulation Power Spectrum procedure (see Elliott & Theunissen, 2009). Additional details regarding the sound editing procedures and acoustic characterization of these vocal excerpts can be found in Arnal et al. (2015).

### Experimental procedure

#### Sound calibration

The intensity of sound presentation was adjusted individually according to the following procedure. Participants were asked to sit comfortably in a chair in a soundproof room, with in-ear earphones (Sennheiser®3.00 Cx) and listened to several presentations of the same sound to determine their hearing threshold. Participants were asked to report when the sound was played by pressing a key. Sounds were played with a random temporal jitter ranging from 3 to 5 seconds. The volume of the sound was set using a self-adjustment procedure increasing twice the volume up to self-reported hearing threshold (starting from -90 digital dB with steps of +/-1.5 digital dB) and decreasing twice the volume down to self-reported hearing threshold (starting from -10 digital dB with steps of +/-1.5 digital dB). The auditory threshold was set as the averaged volume across the four adjustments. The volume of stimulus presentation was then set at 130% of the auditory threshold (for both awake and asleep stimulation).

#### Polysomnographic recording

EEG was recorded with cup electrodes set on Fpz, F3, Fz, F4, T3, C3, Cz, C4, T4, P3, Pz, P4 and Oz positions of the international 10-20 system, and plugged to a 16-electrode EEG amplifier (VAmp model; BrainProducts GmbH). Electrodes were stuck with an adhesive and conductive paste (EC2; Cadwell Industries, Inc.). Additionally, two electro-oculogram (EOG) electrodes were stuck one centimeter above and right to the external right eye canthus and another one centimeter below and left to the external left eye canthus of the participant according to sleep scoring guidelines (Iber et al., 2007). Finally, two electro-myogram (EMG) electrodes were set on the chin, one centimeter of the right and of the left of the dimple of the participant. Electrical signal was recorded at 500 Hz and referenced online to the Fpz electrode. EEG data were filtered online using highpass filter at 0.1 Hz and lowpass filter at 250 Hz.

#### Stimulation protocol

Once the EEG was set up, the participant sat comfortably on a chair in front of a computer, and was asked to stay awake, eyes closed, without moving for one hour, while listening to the two sets of sounds (Fig. 1A). To keep participants fully awake, the experimenters entered the room to check on the participant and explain the instructions once more every 15 min or when signs of drowsiness were detected. The sounds were played one after another with a random jitter of 5 to 11 seconds (drawn from a uniform distribution with a millisecond precision) in alternating blocks of one set of sound or the other. Each block comprised 22 stimuli and lasted about 219 s. Once the awake session was done, the participant was invited to go to bed and fall asleep for the night. The polysomnographic recording of the participant was then monitored online by the experimenters. Overnight stimulation started as soon as the first period of N3 sleep was observed by the experimenter. It was stopped if the participant awoke and resumed when the participant went back to stable (N2, N3 or REM) sleep. To collect comparable amounts of brain responses in wakefulness, NREM and REM sleep (the later not analyzed), each participant was stimulated during the first ∼3 hours of the night - rich in NREM sleep - and during the last expected ∼2 hours of the night - rich in REM sleep (estimated on self-reported sleep schedule). After the night, when the participant was fully awake and ready, a final stimulation session of about 1 hour was performed during wakefulness.

### Sleep scoring and rejection of participants

Sleep was scored by 2 experienced scorers who visually classified the polysomnographic signal (consisting of EEG, EOG and EMG signal) on continuous 20-second epochs according to standards of sleep scoring (Iber et al., 2007). For a detailed summary of sleep measures, see Moyne et al. (2022).

### EEG preprocessing

EEG signal was first epoched from -1 to 2 seconds relative to stimuli onset. Epochs were labeled as wakefulness if the stimulus onset occurred in a 20 s-window of wakefulness, as NREM sleep if the stimulus onset occurred in a 20 s-window of N2 or N3, and as REM sleep if the stimulus onset was in a 20 s-window of REM sleep. Each epoch was then visually inspected and rejected if noisy. Epochs overlapping with micro-arousals (based on manual sleep scoring) were rejected. Then, an Independent Component Analysis (ICA) was computed on all epochs from the two datasets (see section *Stimuli* above). The unmixing matrix was used to compute the source signal from continuous data. Components reflecting eye movements and blinks were rejected and the remaining sources were projected back to the channel space. Channels were then visually inspected and noisy channels were rejected and interpolated from neighboring electrodes. The electrodes were re-referenced to the common average reference.

### Event-Related Potentials

Preprocessed continuous signal was re-segmented into epochs from -1 to 2 s relative to stimuli onsets. Epochs were then low-pass filtered at 30 Hz and baseline corrected using a 500 ms pre-stimulus window.

### Time-frequency analysis

A time-frequency decomposition was carried on continuous preprocessed EEG data. The signal was decomposed in 30 frequencies from 1 to 30 Hz with steps of 1 Hz with Morlet wavelets convolution. Wavelets had a Full Width at Half Maximum (FWHM) linearly decreasing from 1 to 10 cycles of the frequency (1 s for the 1 Hz wavelet and 0.33 s for the 30 Hz wavelet, following the method suggested by Cohen, 2019). Then, the signal was epoched from -2 to 3 s relative to stimuli onset and downsampled by a factor of 10. To compute power changes, the modulus of the complex number was squared. We then took the natural logarithm of this number and multiplied it by 10 (decibel transformation). Finally, a baseline correction was applied to each time series (frequency and trial wise) by subtracting the mean power between -0.5 to 0 s relative to stimulus onset. To compute inter-trial phase coherence (ITPC), the argument function was applied to complex numbers extracted by wavelet convolution to obtain the phase in radian and the following formula was applied:

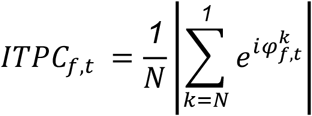

With N the number of trials and 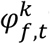 the phase at trial k, at frequency f and time t. Time series (frequency and participant wise) were then corrected by subtracting the average ITPC between -1 and 0 s relative to stimulus onset. For visualization purposes, time-frequency heatmaps of power, ITPC and power modulations related to sound features (pitch and roughness) were upsampled by a factor of 4 in time and frequency by means of cubic spline interpolation. This upsampling procedure was not applied to line plots and statistics.

### Statistics

First-order statistics were computed by averaging trials within units of observation (participant), stimulus type and sleep stage. Exception was made for ITPC, which was calculated on several trials. In this case, the ITPC was calculated for each participant and condition and the result was considered as the first-order statistic. A second exception was made to investigate the linear relation between sigma power and acoustic features. For each participant and in each sleep stage, all stimuli (neutral vocalizations and screams) were included. At each time-point, each frequency and each electrode, the power values across trials were regressed with z-scored roughness values of stimuli in a first regression and with z-scored pitch values in a second regression. Then, the slope of the regression (beta values) was used as first-order statistics for each participant. Finally, second order statistics and p-values were computed across units of observation. Time courses of the paired difference between responses to screams and neutral vocalizations, and influence of roughness and pitch on time courses of the response to all types of vocalizations were statistically tested using cluster permutation. Clusters were selected as consecutive points whose p-values were below 0.05 (*α*_thr_=0.05) as tested with t-tests. Then, cluster values were defined as the sum of the t-values within the cluster (noted as t_cluster_ in the text). 5000 Monte Carlo simulations (N=5000) were performed to compute the distribution of the cluster values across permutations. The p-value of the cluster computed on the distribution of the cluster values is reported in the text as p_cluster_. Statistical testing based on cluster permutation was performed with routines from the Fieldtrip toolbox (Oostenveld et al., 2010). In addition, the original values of power, ITPC, or power gain were averaged across time points of significant clusters; accordingly, the mean across participants and the standard deviation are reported in the text for each significant cluster. In addition, we estimated the effect size of the average values within significant clusters by computing the Cohen’s d as:

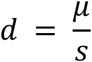

Where *μ* is the above-mentioned mean across participants and s is the above-mentioned standard deviation across participants.

## RESULTS

### Brain responses to screams and neutral vocalizations during wakefulness

We first measured ERPs to scream and neutral conditions. Although stimuli were presented at low sound intensity, auditory ERPs exhibited typical N1-P2-N2 components on Cz (Fig. 2A; see Picton et al., 1974) with expected centro-parietal, central, and fronto-central topographies, respectively. A qualitative comparison suggested that the P2 component was larger for screams than for neutral vocalizations although this difference did not resist cluster-based correction for multiple comparisons (Fig. 2A).

**Figure 2.**
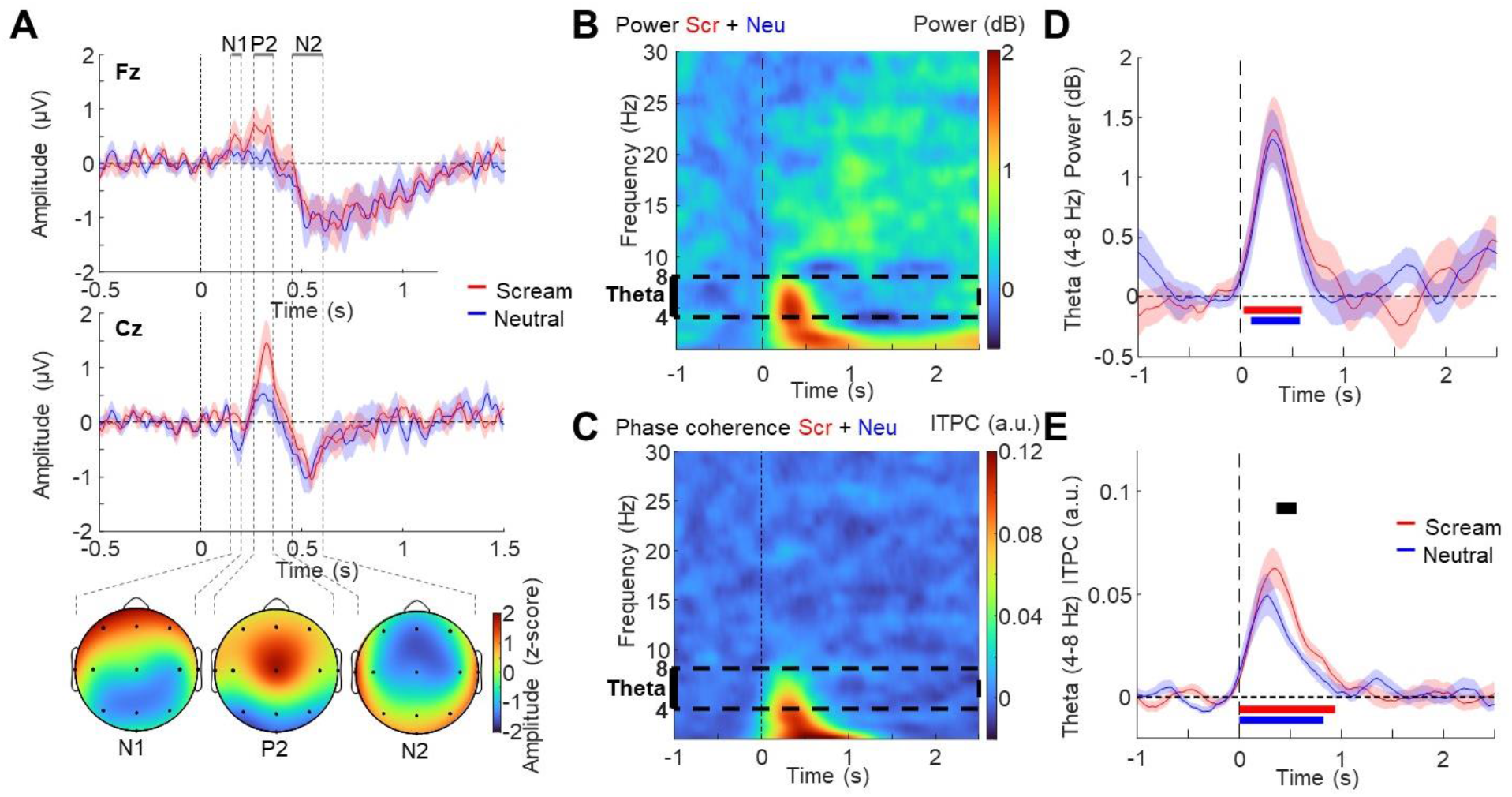
EEG responses to vocalizations during wakefulness. **A**. Screams (red) and neutral vocalizations (blue) evoke typical N1, P2 and N2 ERP components on Fz (top panel), and more prominent responses on Cz (middle panel, and related topographies below). P2 magnitude appears larger for screams than neutral vocalizations although this difference does not resist correction for multiple comparisons. **B**,**C**. The power and ITPC (respectively B and C) time-frequency map of the average responses across all electrodes indicates that vocalizations evoke typical waves in the delta-theta (1–8 Hz) range. To analyze consistently wakefulness and sleep, we excluded the delta frequency range that could potentially be influenced by sleep oscillation (slow-waves). The dashed black rectangle indicates the theta frequency range investigated in further analyses. **D**. Power response was averaged in the theta frequency range for screams (red trace) and for neutral vocalizations (blue trace) but no measurable difference of power between them was observed. **E**. ITPC response was also averaged in the theta range and shows that auditory phase-locked responses are larger for screams (red trace) than for neutral vocalizations (blue trace). Shaded surfaces in plots indicate SEM. Significant statistical differences (corrected using cluster-based permutations) are plotted as thick horizontal lines (red for screams against 0; blue for neutral vocalizations against 0 and black for the difference screams vs neutral vocalizations).

We then used time-frequency resolved analyses to investigate potential differences in amplitude or time-consistency across single-trial EEG responses to screams and neutral vocalizations. The time-frequency map of the averaged response in scream and neutral conditions showed an overall increase of power (Fig. 2B) and intertrial phase coherence (ITPC; Fig. 2C) in the theta and delta frequency ranges over the time of stimulation. Focusing on auditory responses in the theta frequency-range, we found that both screams and neutral vocalizations induced an increase of power (Fig. 2D; screams: significant cluster from 0.020 to 0.600 s, mean power difference±SD (dB): 3.12e-01±5.63e-01, N=13, t_cluster_=126.91, p_cluster_=0.003, d=0.55; neutral: significant cluster from 0.100 to 0.580 s, mean power difference±SD (dB): 3.05e-01±4.57e-01, N=13, t_cluster_=109.36, p_cluster_=0.007, d=0.67). However, we did not observe a significant theta power difference between the two conditions (Fig. 2D). We then measured the time-alignment of brain response and found that the increase in ITPC was significant for screams (Fig. 2E; significant cluster from 0 to 0.940 s; mean ITPC difference±SD (a.u.): 3.53e-02±1.57e-02, N=13, t=273.27, p<0.001, d=2.25) and for neutral vocalizations (significant cluster from 0 to 0.820 s; mean ITPC difference±SD (a.u.): 2.87e-02±1.79e-02, N=13, t=212.40, p<0.001, d=1.61). We also found a larger increase of ITPC for screams compared to neutral vocalizations between 0.360 and 0.560 s post-stimulus onset (Fig. 2E; significant cluster from 0.360 to 0.560 s; mean ITPC difference±SD: 1.98e-02±2.29e-02 a.u., N=13, t_cluster_=32.00, p_cluster_=0.046, d=0.86), showing that roughness enhanced phase-concentration in the theta range across trials.

### Brain responses to screams and neutral vocalization during NREM sleep

We then wondered whether, similarly to wakefulness, brain responses to screams are more consistent in time in NREM sleep. We first compared ERPs to screams and neutral vocalizations during NREM sleep. Consistent with the literature, both conditions evoked large deflections (slow-waves) with highest amplitude on frontal electrodes (Fig. 3A; top panel) and “earlier” components (before 600 ms) on the central Cz electrode (Fig. 3A; bottom panel). The three main components of evoked slow-waves (P200, N550 and P900) typically displayed highest amplitudes on frontal electrodes. We did not find a significant difference between screams and neutral vocalizations in the time courses of ERPs on Fz or Cz electrodes.

**Figure 3.**
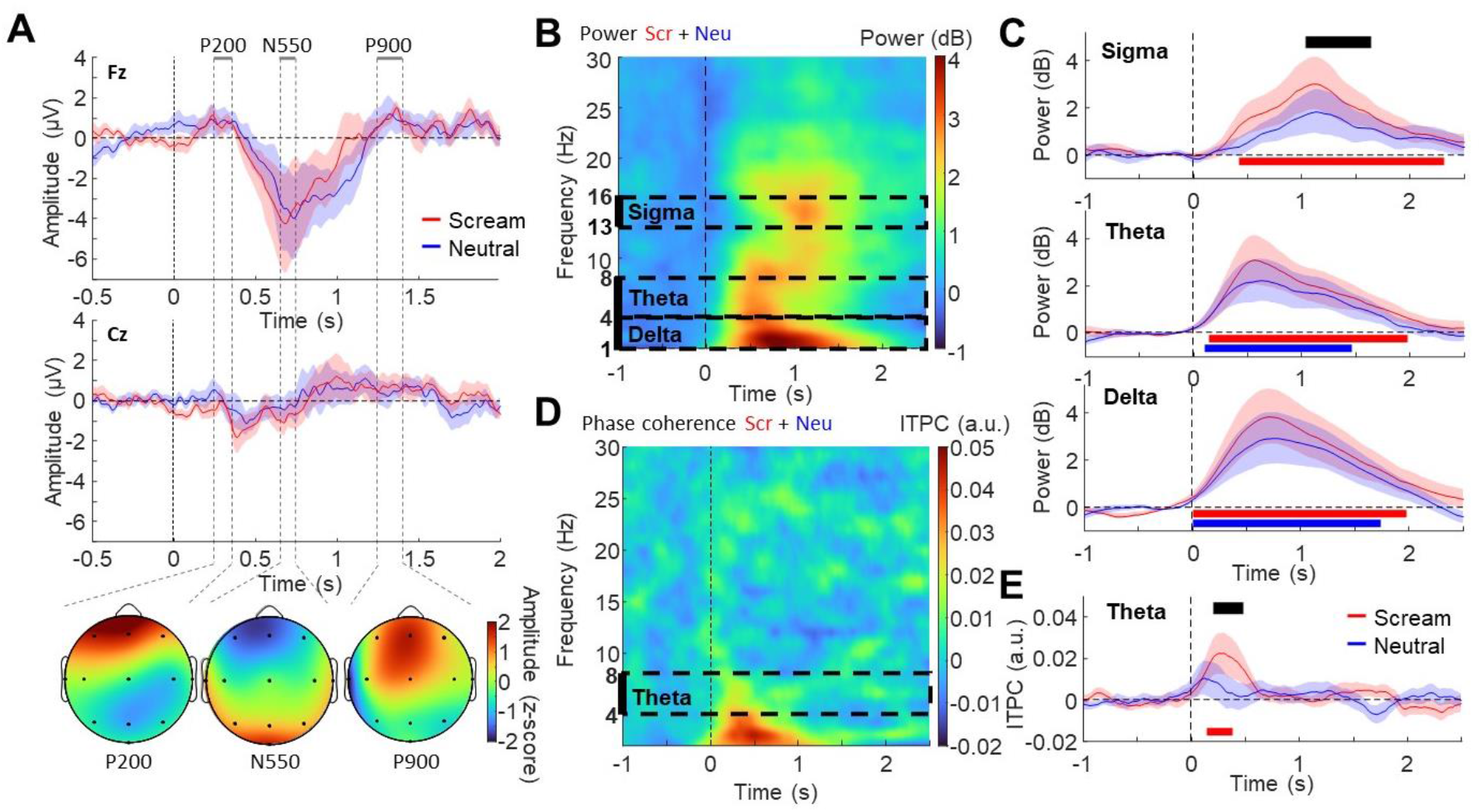
EEG responses to vocalizations during NREM sleep. **A**. Screams (in red) and neutral vocalizations (in blue) evoke slow-waves, with maximal amplitude on Fz electrode (top panel) and less prominent on Cz (middle panel) although initial auditory-evoked potentials can be visually observed on Cz before 0.6 s. Topographies (at the bottom) of the P200, N550 and P900 are typical of evoked slow-waves during sleep, with maximal amplitude on frontal electrodes. **B**,**C**. ITPC and power (respectively B and C) time-frequency map averaging responses across all electrodes indicates that, as during wakefulness, vocalizations evoke phase-locked responses in the delta-theta (1–8 Hz) range. The power time-frequency map also suggests an increase of waves in the sigma frequency range. Dashed black rectangles highlight the frequency ranges (delta, theta and sigma) explored in further analyses. **D**. Power responses averaged in the sigma (top panel), theta (middle panel) and delta (bottom panel) frequency ranges reveal an increase in all ranges investigated for both screams and neutral vocalizations except in the sigma range where only screams induce a significant increase. **E**. Averaged ITPC response in the theta (4–8 Hz) range are larger for screams (red trace) than for neutral vocalizations (blue trace), which are merely detectable. Plain lines indicate the mean response across participants and shaded surfaces the SEM in plots of D and E. Significant statistical differences (corrected using cluster-based permutations) are plotted as thick horizontal lines (red for screams against 0; blue for neutral vocalizations against 0 and black for the difference screams vs neutral vocalizations).

Then, we evaluated whether the power of evoked oscillations that are typical of ERPs (i.e. theta frequency range) and NREM sleep (i.e. spindles in the sigma frequency range and slow-waves in the delta frequency range) is influenced by the type of vocalization. Using the same time-frequency decomposition approach as in the wake data, we found that spectro-temporal response patterns were mostly conserved during sleep. We first observed that screams and neutral vocalizations evoked low-frequency responses in the delta [1-4 Hz] frequency range (Fig. 3B; bottom right panel; significant cluster of the difference screams vs baseline from 0 to 1.980 s; mean power difference±SD (dB): 2.06±2.40, N=13, t_cluster_=295.43, p_cluster_=0.003, d=0.86; significant cluster of the difference neutral vocalizations vs baseline from 0 to 1.740 s; mean power difference±SD (dB): 1.56±2.10, N=13, t_cluster_=238.90, p_cluster_=0.014, d=0.74), as well as in the theta [4-8 Hz] frequency range (Fig. 3B; middle right panel; significant cluster of the difference screams vs baseline from 0.140 to 1.980 s; mean power difference±SD (dB): 1.46±1.70, N=13, t_cluster_=265.62, p_cluster_=0.003, d=0.86; significant cluster of the difference neutral vocalizations vs baseline from 0.100 to 1.460 s; mean power difference±SD (dB): 1.08±1.66, N=13, t_cluster_=175.50, p_cluster_=0.020, d=0.65). Focusing on the theta [4–8 Hz] frequency range, we found an overall increase of auditory responses in sleep relative to wakefulness at the single-trial level in all conditions (Supplementary Fig 1A; Significant cluster from 0.520 to 1.500 s; Mean power difference±SD (dB): -1.78±2.42, N=13, t=-157.27, p=0.005, d=0.73) and an overall decrease of ITPC (Supplementary Fig 1B; Significant cluster of the difference wake vs NREM sleep from 0.100 to 0.620 s; Mean ITPC difference±SD (a.u.): 1.62e-02±2.61e-02, N=13, t=80.85, p=0.007, d=0.62). When splitting across auditory conditions, early responses to screams in the theta range exhibited significant alignment (Fig. 3C; right panel; significant cluster from 0.140 to 0.380 s; mean ITPC difference±SD (a.u.): 2.04e-02±3.12e-02, N=13, t_cluster_=30.16, p_cluster_=0.039, d=0.65) - although partially reduced compared to wakefulness - whereas neutral vocalizations did not modulate theta ITPC (no statistical difference of the ITPC vs baseline). Moreover, ITPC magnitude was larger for screams than for neutral vocalizations in a similar time window [0.200 to 0.480 s] as observed during wakefulness (significant cluster from 0.200 to 0.480 s; mean ITPC difference±SD: 1.54e-02±2.04e-02 a.u., N=13, t_cluster_=38.93, p_cluster_=0.031, d=0.75). This result suggests that roughness, even at a faint intensity, promotes fast and consistent processing of auditory stimuli, in line with our main hypothesis.

In addition to an increase of theta power and slow-waves, time-frequency analyses revealed an increase in higher frequencies between 13 and 16 Hz, a range that is typical of fast spindles (Fig. 3B; left panel; see Fernandez & Lüthi, 2019). As stimulus salience generally increases sleep spindle generation (Blume et al., 2017, 2018), we tested whether high-sigma power increase was influenced by the type of vocalization. First, we found that sigma power increased in response to screams (significant cluster from 0.420 to 2.320 s; mean power difference±SD (dB): 1.48±1.93, N=13, t_cluster_=254.22, p_cluster_=0.002, d=0.76) but not after neutral vocalizations. Comparing responses between conditions, we found that screams induced larger responses in the sigma range at late latencies, between 1.040 and 1.640 s (significant cluster from 1.040 to 1.640 s; mean power difference±SD: 9.20e-01±1.23 dB, N=13, t_cluster_=76.41, p_cluster_=0.011, d=0.75; Fig. 3B; top right panel).

### Relationship between roughness and evoked sleep oscillations

To characterize whether pitch and roughness are differentially processed during NREM sleep, we tested how each of these acoustic features affected the generation of slow-waves and spindles. To this end, we used regression analyses between sound roughness or pitch and the power of frequency resolved responses. As expected from our previous observations, a qualitative inspection of these maps suggested that roughness (Fig. 4A), unlike pitch (Fig. 4B), enhanced the power of brain responses in the delta and sigma frequency ranges (dashed black rectangles), consistent with an increased generation of slow-waves and spindles by roughness, respectively. We then focused our analyses on the specific frequency-bands of slow-waves and spindles, which are typical spectral signatures of exogenous processing during sleep. To do so, we statistically tested the coefficients of the regressions with roughness and pitch across time in the high sigma(13-16 Hz; Fig. 4C) and delta (1-4 Hz; Fig. 4D) range. In the delta range, this analysis revealed that roughness –but not pitch– enhanced the amplitude of slow-waves, yielding a significant cluster between 0.500 to 0.940 s (mean power gain±SD: 3.78e-01±5.50e-01 dB, N=13, t_cluster_=56.28, p_cluster_=0.036, d=0.69; Fig. 4D). Consistent with the typical topographic expression of slow waves, the increase of delta power with roughness was maximal on Fz (Fig. 4F). Using a similar approach, we then measured the effects of roughness and pitch features on the generation of spindles. This analysis showed a significant relationship between roughness and sigma power in a later 0.940 to 1.300 s time-window (mean power gain±SD: 4.90e-01±7.53e-01 dB, N=13, t_cluster_=43.73, p_cluster_=0.046, d=0.65; Fig. 4C), whereas no significant power gain was observed in the regression with pitch values. The effects of roughness displayed a typical topography of spindles, with a larger increase on central electrodes (Fig. 4E).

**Figure 4.**
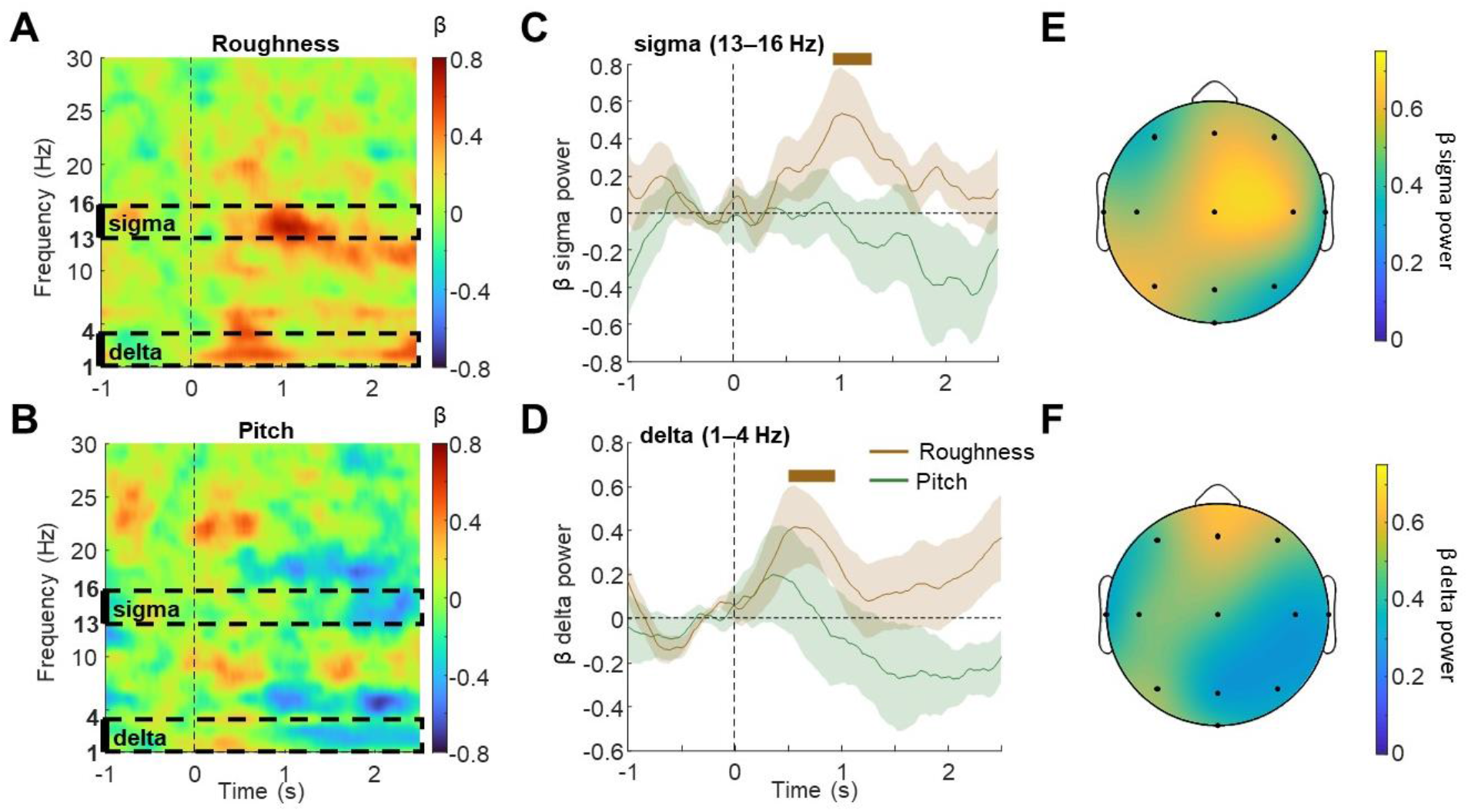
Roughness (but not pitch) enhances evoked responses during sleep. **A**,**B**. Regression of evoked responses (at each time point and frequency) by stimulus roughness (A) or pitch (B). Black dotted rectangles indicate time-frequency windows of interest in sleep-related frequency ranges, namely in the sigma (13–16 Hz) and delta (1–4 Hz) ranges, for subsequent statistical analyses. **C**,**D**. Time-course of sigma (C) and delta (D) power regressions by roughness (brown) and pitch (green). While the pitch of vocalizations does not seem to influence brain responses during sleep, roughness induces a significant delta-followed by a sigma-power increase during auditory stimulation. Shaded surfaces indicate SEM. **E**,**F**. Topographies of sigma (E) and delta (F) responses induced by roughness, averaged across time- and frequency-window of significance (thick black lines in C and D). The spatial and frequency selectivity of these topographies suggests that roughness enhances typical sleep-related cerebral responses, namely sleep spindles (central electrodes) and slow-waves (frontal electrodes).

## DISCUSSION

In the last decades, a large body of research has supported the idea that the brain maintains a certain level of responsiveness toward environmental events during sleep (Beh & Barratt, 1965; Blume et al., 2017, 2018; Chen et al., 2016; Formby, 1967; Hayat et al., 2022; Legendre et al., 2019; Moyne et al., 2022). However, sensory stimuli often do not reach conscious perception during sleep and whether the brain prioritizes certain acoustic features over others remains unclear. Sound loudness, which indexes the spatial distance and physical force of the source, is perhaps the most relevant feature to process during sleep. On the other hand, whether we are more sensitive to certain spectral features than others at low sound intensity during sleep is unknown. We assumed that, owing to their aversiveness and their propensity to capture attention, temporally salient, rough sounds (as featured in human screams or wakening devices such as alarm clocks, buzzers) are likely to be processed more efficiently than smoother neutral sounds during sleep (Arnal et al., 2015, 2019).

### Reliable brain responses to screams during NREM sleep

Here, we show that, even at a low intensity, screams induced more reliable, better time-locked brain responses than neutral vocalizations across both wakefulness and NREM sleep. This was expressed as increased ITPC of evoked responses in the theta frequency range. The ITPC metrics can be slightly influenced by the amplitude of waves but mainly reflects the degree of phase-consistency across trials of stimuli-evoked EEG waves (van Diepen & Mazaheri, 2018). As our power analysis did not reveal a difference of power induced by our two types of stimuli, it is likely that the difference of ITPC reflects a better alignment in time of evoked brain responses across trials. Of note, studies using an auditory perceptual learning paradigm found that ITPC, but not power, of evoked responses indexed the detection of hard-to-notice target sounds in wakefulness and sleep (Andrillon et al., 2015, 2017). The authors suggested that such ITPC increase reflects the formation of reliable brain responses through perceptual learning and the clear detection of auditory patterns. Similarly, scream’s acoustic features might promote their detection, inducing more reliable brain responses and increasing ITPC by enhancing the temporal alignment and synchrony of neural populations across emotional and auditory areas. This is consistent with recent findings that rough sounds enhance neural synchronization throughout temporal cortical and subcortical limbic structures (Arnal et al., 2019). In that study, evoked high-gamma power varied linearly with the frequency of click trains (i.e., a type of high roughness sound composed of series of *click*), suggesting a linear relationship between response amplitude of neural populations and sound energy in auditory cortical areas. On the other hand, brain responses aligned maximally with stimuli in the roughness range (∼30–80Hz), supporting the current finding that stimuli in this range enhance neural synchronization. Noteworthy, although it is moderately diminished compared to wakefulness, auditory steady-state response and ITPC in the delta and theta frequency range following 40Hz click trains can still be observed in NREM sleep (Hayat et al., 2022; Lustenberger et al., 2018; Picton et al., 2003). In the visual domain, emotional stimuli, like angry or happy faces, can boost the activity of neural populations in emotional (e.g. amygdala) and sensory cortices (Domínguez-Borràs et al., 2019; Guex et al., 2020; Vuilleumier et al., 2001). In turn, emotional items are detected faster than neutral items (Eastwood et al., 2001; Öhman et al., 2001). Some features of emotional prosody of meaningless utterances such as laughs, cries, and screams, lead to higher metabolic activity of the amygdala and enhanced auditory processing (Arnal et al., 2015; Frühholz et al., 2012; Frühholz & Grandjean, 2013; Grandjean et al., 2005; Sander & Scheich, 2001). Thus, roughness enhances screams processing by exogenously entraining neuronal synchrony in emotion-related brain regions and attentional networks.

How rough sounds target brain networks involved in attentional and affective processing remains a matter of active research (Arnal et al., 2015, 2019; Farahani et al., 2021). In line with recent evidence, the current results support the hypothesis that rough sounds promote aversion by triggering sustained neuronal synchronization across widespread subcortical and cortico-limbic networks involved in exogenous attentional processing (Arnal et al., 2019). The spatio-temporal profile of such responses is compatible with the recruitment of non-canonical auditory pathways ultimately targeting salience processing systems (Arnal et al., 2019; Farahani et al., 2021). The precise neural routes underlying such extensive synchronization by rough sounds remain unclear. However, that roughness processing appears selectively preserved during sleep raises interesting questions. One hypothesis, supported by recent animal work (Kim et al., 2015), points towards a potential role of deep cerebral circuits involved in sleep regulation. This idea is also supported by experimental observations that brain responses to sounds in the roughness range are reduced in comatose patients with suspected alteration in the thalamus and brainstem (Firsching et al., 1987). The larger evoked slow-waves and spindles in response to screams are also consistent with a potential involvement of arousal-promoting neuromodulatory groups of the brainstem (Jahnke et al., 2012; Siclari et al., 2014). By targeting and synchronizing such systems during sleep, rough sounds might more efficiently arouse the cerebral cortex to awaken the brain and promote faster reactions to signaled danger. However, although fascinating, the idea that rough sounds selectively recruit such pathways remains largely hypothetical and needs to be formally traced anatomically and functionally using animal models.

Owing to its capacity to induce sustained and reliable neuronal synchrony in attentional networks, roughness appears a powerful way of communicating danger. By triggering reliable responses in a state of reversible unresponsiveness such as NREM sleep, roughness constitutes a particularly efficient and adaptive signal. The current findings suggest that such features remain efficiently processed even at very low intensity during sleep. This ensures that warning signals such as screams can be optimally processed from a distance by sleeping conspecifics.

### Acoustic roughness boosts the generation of sleep spindle and slow-waves

By stimulating participants with screams and neutral vocalizations during sleep, we showed that screams yield a transient increase in sigma power, as compared to neutral vocalizations. In addition, we observed that the rougher the vocalization, the stronger the evoked sigma power was. Sleep spindles, which are the main source of sigma power during NREM sleep, are linked to sleep maintenance, as they are more abundant in the nights of people with stable sleep (Dang-Vu et al., 2010) and during periods of sleep more resistant to disturbances (Lecci et al., 2017). In addition, sleep spindles can be triggered by external stimulations and are potentiated by stimulation salience (Blume et al., 2017, 2018; Sato et al., 2007). Hence, some authors proposed that evoked slow-waves and sleep spindles represent intermediate responses on the arousal spectrum between small brain responses such as ERPs and complete micro-arousal or awakening (Andrillon & Kouider, 2020; Blume et al., 2018; Schabus et al., 2012). Thus, the stimulus would be salient enough to activate broad networks but would reflect such a small threat (in our case, the low intensity might dampen the threatening aspect of screamed vocalizations) that awakening would not be necessary. To revert to quiet sleep, spindle generation would prevent further processing of sensory stimulation.

Vocalizations’ roughness also increased early delta power, a marker of slow-wave activity. Much like what was reported above for spindles, the generation of slow-waves appears to be linked to the salience of stimuli (Halász, 2005). Thus, as rough vocalizations evoked more slow-waves and spindles in the present study, we argue that roughness, even when presented at very low intensity, constitutes an efficient attribute to mobilize exogenous attentional systems during wakefulness as well as during sleep. Whether roughness increases the probability to generate spindles and slow-waves or whether it increases their intensity (larger spindles and slow-waves) cannot be directly answered by our analyses. Yet, phase consistency analysis revealed that initial responses evoked in the theta range suggests that brain responses to screams are more reliable across repetitions. Thus, spindles and slow-waves might also be more systematically generated.

This research highlights two crucial findings: first, the direct comparison between the evoked responses to screams and neutral vocalizations in NREM sleep supports that 1) roughness is a salient acoustic feature irrespective of vigilance state and orthogonal to sound intensity and 2) sigma power is a reliable marker of roughness-related stimulus salience in NREM sleep. These findings might account for the aversiveness of rough snoring sounds and their detrimental effects on sleep (Fischer et al., 2016). Many human activities generate rough sounds (Arnal et al., 2015; Raggam et al., 2007), which might enhance sleep disturbance by enhancing the recruitment of attentional and emotional brain circuits sleep (Eberhardt, 1988; Muzet, 2007). These results further highlight the importance of considering specific sound attributes – beyond intensity– such as roughness as a relevant factor to promote healthier sleeping environments.

## ACKNOWLEDGMENTS

This work was supported by the National Center of Competence in Research (NCCR) Affective Sciences (financed by the Swiss National Science Foundation, 51NF40-104897; and hosted by the University of Geneva), and by grants from the Swiss National Science Foundation (320030-159862 to S.S.; Ambizione - PZ00P3-148112 to J.D.B.) and the Fondation Pour l’Audition FPA RD-2020-10 (L.H.A.) The authors thank Dr. Keith B. Doelling for helpful comments on the manuscript.

