## Supplementary material for "Scream’s roughness confers a privileged access to the brain during sleep"

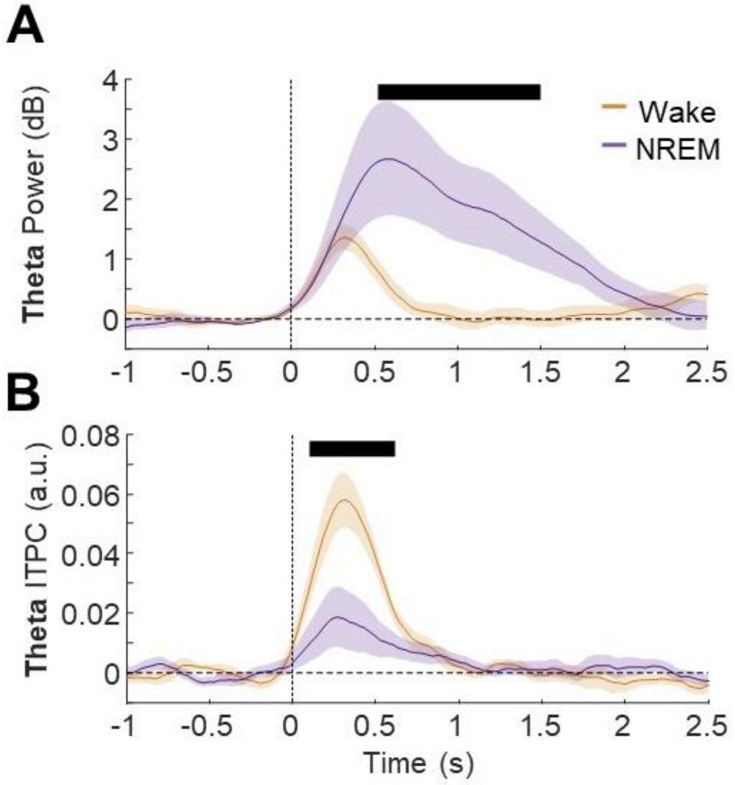


**Supplementary Figure 1**. **A,B.** Comparison of power (A) and ITPC (B) of brain responses at wakefulness (orange) and in NREM (dark blue) in the theta frequency range to screams and neutral vocalizations together. Plain line represent the mean across participants and shaded area the SEM.
